# Integrating Shear Flow and Trypsin Treatment to Assess Cell Adhesion Strength

**DOI:** 10.1101/2023.09.26.559598

**Authors:** Antra Patel, Bhavana Bhavanam, Trevor Keenan, Venkat Maruthamuthu

**Affiliations:** Department of Biological Sciences, Old Dominion University, Norfolk, VA 23529 USA; Department of Mechanical & Aerospace Engineering, Old Dominion University, Norfolk, VA 23529 USA

**Author notes:** Corresponding author’s address: Venkat Maruthamuthu, Mechanical & Aerospace Engineering, Engineering Systems Building 2123J, 1 Old Dominion University, Norfolk VA 23529, United States.

**Keywords:** cell adhesion, microfluidics, shear flow, trypsin, adhesion strength

## Abstract

Cell adhesion is of fundamental importance in cell and tissue organization, and for designing cell-laden constructs for tissue engineering. Prior methods to assess cell adhesion strength for strongly adherent cells using hydrodynamic shear flow either involved the use of specialized flow devices to generate high shear stress or used simpler implementations like larger height parallel plate chambers that enable multi-hour cell culture but generate low shear stress and are hence more applicable for weakly adherent cells. Here, we propose a shear flow assay for adhesion strength assessment of strongly adherent cells that employs off-the-shelf parallel plate chambers for shear flow as well as simultaneous trypsin treatment to tune down the adhesion strength of cells. We implement the assay with a strongly adherent cell type and show that shear stress in the 0.07 to 7 Pa range is sufficient to dislodge the cells with simultaneous trypsin treatment. Imaging of cells over a square centimeter area allows cell morphological analysis of hundreds of cells. We show that the cell area of cells that are dislodged, on average, does not monotonically increase with shear stress at the higher end of shear stresses used and suggest that this can be explained by the likely higher resistance of high circularity cells to trypsin digestion. The adhesion strength assay proposed can be easily adapted by labs to assess the adhesion strength of both weakly and strongly adherent cell types and has the potential to be adapted for substrate stiffness-dependent adhesion strength assessment in mechanobiology studies.

## Introduction

Most metazoan cells adhere to their surroundings using specific proteins embedded in their cell membrane [1]. For instance, mammalian cells in solid tissues such as epithelia typically adhere to the extra-cellular matrix (ECM) beneath as well as cell-cell adhesion molecules laterally [2]. The level of adhesion at each interface and via each specific family of receptors is functionally important in a myriad of ways, influencing cell survival [3], proliferation [4], growth [5] and migration [6]. It is also an important consideration in the design of tissue constructs and organoids [7]. The level of cell adhesion between a cell and its partner surface depends on many factors, including the level of expression of the particular cell adhesion receptor, the level of contractility of the cell and the geometry of the contact. It also depends on more involved factors such as the cell signaling state [8-12] that in turn influences the adhesive state of the adhesion proteins as well as the organization of the cell cytoskeleton that supports adhesion [13]. The level of adhesion is also influenced by the stiffness of the surface to which the cell is adhering [14, 15]. The level of cell adhesion can be mechanically characterized using the cell adhesion strength, which is the force required to disrupt the adhesion between a cell and a surface to which it adheres [16]. Cell adhesion strength is a biophysical parameter relevant to understanding cell behaviors in various biological contexts [17], including as a potential marker for metastatic cancer cells [18].

To date, many assays have been proposed and employed in the measurement of cell adhesion strength [19]. While most such assays, including the one proposed here, are demonstrated using cells adherent on ECM-coated surfaces, these assays are amenable to use with many adhesion systems, including cell-cell adhesion systems, with a judicious choice of coating of specific proteins on the surface to which the cells adhere [15, 20]. Cell adhesion strength assays can be broadly classified into single-cell methods, which involve probing of cells one at a time, and bulk assays that can assess the adhesion strength of many cells in each assay [19]. Bulk adhesion strength assays, by definition, yield data on a larger number of cells and can aid in capturing the heterogeneity among cells. For this reason, we will focus on these bulk assays here. These assays include simple mechanical disruption of cell collectives by pipetting [21], centrifuge assays (which involve spinning of surfaces with adherent cells, with the surface normal perpendicular to the spinning axis) wherein cells are detached by centrifugal forces [22], spinning of surfaces with adherent cells (with the surface normal parallel to the spinning axis) wherein the cells are detached by hydrodynamic shear forces [23, 24], radial [25, 26], cone and plate [27], jet impingement [28] and parallel plate (including microfluidic) flow methods [29] wherein the cells are detached by hydrodynamic shear forces. Note that most of the bulk adhesion strength assays do permit the acquisition of cell-level data, where characteristics of individual cells can still be obtained.

Parallel plate assay configurations [29-33] typically involve chamber heights >200 μm [19] and can produce shear stress of up to a couple of tens of Pa. Microfluidic shear flow chambers are also effectively parallel-plate chambers but typically involve chamber heights <100 μm [19] and can produce up to 100s of Pa shear stress. However, the construction of such microfluidic chambers often involves the use of photolithography [34], which may not be accessible to all labs. Use of ‘off-the-shelf’ parallel-plate chambers that can either be procured as such or constructed more readily and easily can benefit widespread use of shear flow for adhesion strength assessment. But the shear stress produced by these chambers with larger heights is not sufficient to dislodge strongly adherent cells. The required higher shear stress was realized using small height chambers previously [35, 36]. Furthermore, some of the shear flow studies using parallel plate chambers or glass capillaries assessed the adhesion strength of cells that were plated for only an hour or less [20, 33]. Despite rapid area expansion over the first 10 min [37], adherent cell types typically take more than an hour to fully spread and adhere to the surface on which they are plated, with the final cell area sometimes attained only at around two hours after cell plating [38, 39]. Thus, plating of cells for more than an hour, preferably 2 h or more before the assay is desirable. Custom-made microfluidic channels that employ photolithography enable long-term cell plating via more involved channel designs that alleviate nutrient mass transfer limitations [34, 36]. But since realization of these more involved microfluidic designs is not practical for all labs, a more accessible approach is required.

Here, we sought to propose and demonstrate a modified parallel plate shear flow assay that can employ off-the-shelf parallel plate flow chambers for assessing cell adhesion strength. Since these chamber heights are usually greater than 100 μm, we sought to simultaneously include another factor – trypsin digestion of cell adhesion proteins at the cell-surface interface – that can aid in adhesion strength assessment. Trypsin is a serine protease that is near-universally used to detach cells from culture substrates during routine cell culture and passaging. Depending on the concentration of trypsin used [40], trypsin can detach cells on the timescale of minutes to tens of minutes. We thus propose and demonstrate the simultaneous employment of shear flow and trypsin treatment to assess the cell adhesion strength of a strongly adherent cell type and elucidate the type of data that can be obtained from such an assay.

## Materials and Methods

### Cell culture and reagents

Madin-Darby Canine Kidney (MDCK II) cells were grown in DMEM (Dulbecco’s modified Eagle’s medium, Corning Inc., Corning, NY) supplemented with 10% Fetal Bovine Serum (FBS) (Corning Inc., Corning, NY), sodium pyruvate, L-Glutamine and 1% Penicillin/Streptomycin at 37 °C under 5% CO_2_. For shear flow experiments, cells were plated in an off-the-shelf parallel plate flow chamber (details in the following sub-section) with a standard glass microscope slide as base. The bottom glass surface of the chamber was first coated with the extra-cellular matrix collagen I as follows: the chamber was filled with 0.2 mg/mL collagen I in 0.1 M acetic acid, allowed to incubate for 15 min, and then washed with phosphate buffered saline (PBS) thrice. MDCK cells were non-enzymatically removed from culture dishes (0.5 mM Ethylenediaminetetraacetic (EDTA), as in Versene, Thermo Fisher Scientific, Waltham, MA) and plated into the microfluidic chamber by pipetting in through the inlet. Cells were allowed to adhere for 2 hours. Hoechst 33342 (diluted from a 10 mg/mL stock solution in PBS and used at 1:2000 for 15 min with cells) was used as a live stain for the nucleus and CellBrite Red membrane dye (diluted in PBS and used at 1:200 for 15 min with cells) was used as a live stain for the plasma membrane.

### Shear flow set-up

The microfluidic chamber used for shear flow experiments was off-the-shelf (Ibidi, Fitchburg, WI), with a nominal height of 150 μm and width of 5 mm. The liquid medium (trypsin solution) used for the shear flow experiments was a Hank’s balanced salt solution with 0.25% trypsin, 0.1% EDTA and no calcium or magnesium (Corning Inc., Corning, NY). Flow was driven from a plastic syringe with a 50 mL nominal capacity (BD, Franklin Lakes, NJ) using a digital syringe pump (PHD 2000, Harvard Apparatus, Holliston, MA).

### Imaging

Imaging was performed using a Leica DMi8 epifluorescence microscope (Leica Microsystems, Buffalo Grove, IL) with a Clara cooled CCD camera (Andor Technology, Belfast, Ulster, UK) and an airstream incubator (Nevtek, Williamsville, VA). Hoechst 33342 and CellBrite Red were imaged via their respective fluorescence channels. Cell images (phase and the above mentioned two fluorescence channels) were acquired over a ∼30 mm x ∼3 mm rectangular area by using the tile scan feature of the LAS X software (Leica Microsystems, Buffalo Grove, IL). This involved frame by frame acquisition (36 × 5 = 180 frames) using a 10x objective magnification followed by image stitching, leading to a large ∼47000 pixel x ∼4900 pixel composite image for each experiment. Note that one can also use the Image Stitching plugin in Fiji, an open source image processing package based on ImageJ, or a function such as ImageStitch in Mathematica (Wolfram Research) to accomplish the stitching of many partially overlapping images acquired over a large area. At time zero, phase images, nuclear fluorescence images and plasma membrane fluorescence images were acquired. For subsequent time points, only the nuclear fluorescence images were acquired.

### Data analysis

For analysis, we first identified cells that are single (i.e., not adherent to any other cell) and are at least one cell diameter away from neighboring cells (so that shear flow and trypsin treatment, rather than inter-cellular mechanical interactions [41, 42], determine their coming off the surface in the experiment). Using image stacks of nuclear fluorescence images (acquired at sequential time points), we used ImageJ to identify the time point at which each of these single cells came off the surface (as revealed by their absence in the composite image of all fluorescent nuclei at that time point, but not the immediately preceding time point). The plasma membrane images as well as the phase images of the cells were used to manually segment each cell using ImageJ (NIH, Bethesda, MD). ImageJ was also used to extract the shape characteristics of the segmented cells – the cell area and the circularity, defined as 4π area/(perimeter^2^). The mean of cell areas or circularities of cells that were dislodged in adjacent shear stress categories (low and medium or medium and high) were compared using a ttest, with a Bonferroni correction. ** indicates p < 0.01 and *** indicates p < 0.001.

## Results and Discussion

Given the need to consider cells that have been cultured for at least an hour or two to allow full cell spreading and adherence, we wanted to develop an adhesion strength assay based on shear flow but with an off-the-shelf chamber that enables multi-hour cell culture before the assay. As outlined in Fig.1, the adhesion strength of adherent cells like the epithelial cell line MDCK or fibroblast 3T3 [34] is such that high shear forces are required to rupture the cell’s adhesion with an extra-cellular matrix (ECM) coated surface. This necessitates microfluidic chambers with a small height, which translates to high wall shear stress for a given flow rate (Fig. 1). The major issue with culturing cells in such highly confined chambers (before the assay) is the adverse impact of cell nutrient mass transfer limitations on cell viability. This in turn reduces the allowable time for the cells to adhere after plating in such a chamber [33]. While some chambers avoid this limitation with more intricate designs [34, 36], we wanted to employ off-the-shelf chambers that are more amenable for wider use in labs. Since such off-the-shelf chambers have greater heights, this limits the maximum wall shear stress that can be realized (Fig. 1). Thus, we chose to use the enzyme trypsin to tune down the adhesion strength (via cleavage of a fraction of cell adhesion receptors) such that lower shear forces can dislodge the cells (Fig. 1).

**Figure 1.**
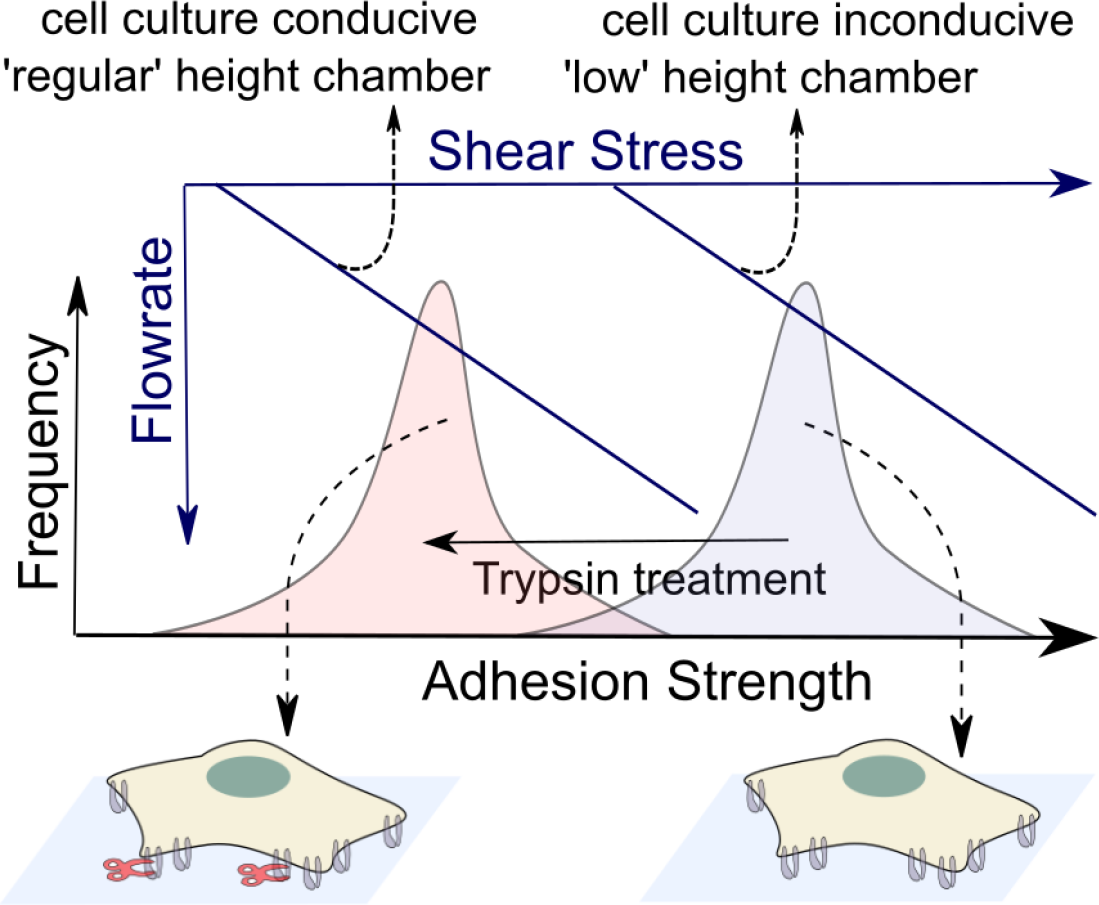
Schematic depiction of the rationale behind our proposed method to assess the adhesion strength of adherent cells. There is a distribution (grey bell-shaped curve) of adhesion strengths for adherent cells (schematically shown at the bottom right) under normal conditions. The shear stress required to probe these adhesion strengths in a flow-based assay is correspondingly high. Under trypsin treatment, the adhesion strengths are lower in magnitude (light red bell-shaped curve) due to some of the adhesions being removed by trypsin digestion (schematically shown at the bottom left). The shear stress required to probe these lower resultant adhesion strengths in a flow-based assay is correspondingly lower.

Fig. 2A shows a schematic of our setup, where a syringe pump is used to flow aqueous media containing trypsin to enable simultaneous shear flow and trypsin treatment in order to assess the cell adhesion strength. We employed an off-the-shelf parallel plate chamber of rectangular cross-section, with width w = 5 mm and height h = 164±5 μm, as determined by imaging the top and bottom walls of the channel using a light microscope. Thus, w/h ∼30 and the hydraulic diameter D_*h*_, given by equation (1) below is computed to be ∼ 320 μm for our channel.

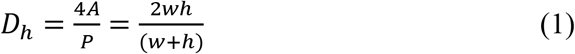

where A is the cross-sectional area and P the perimeter of the flow channel. The Reynolds number *Re*, given below is less than 100 for all flowrates used in our study (Fig. 2B).

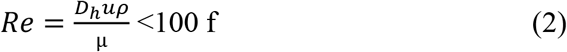

with u, the average flow velocity, ρ, the density of trypsin solution similar to that of water ∼1000 kg/m^3^, μ, the viscosity of the trypsin solution similar to that of phosphate buffered saline [43], ∼0.9 mPa.s at room temperature. The maximum volumetric flowrate Q employed by us was 10 mL/min (Fig. 2B). Thus, the entrance length [44] L_*e*_, given below

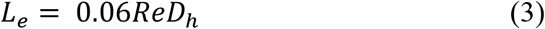

was ∼ 1.5 mm. Therefore, we chose to image the cells in the chamber at a distance significantly greater (∼10 mm) than the entrance length from the fluid inlet. For laminar flow in rectangular channels with a large w/h, the friction factor *f* is given by

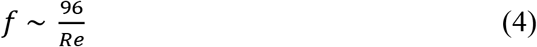

**Figure 2.**
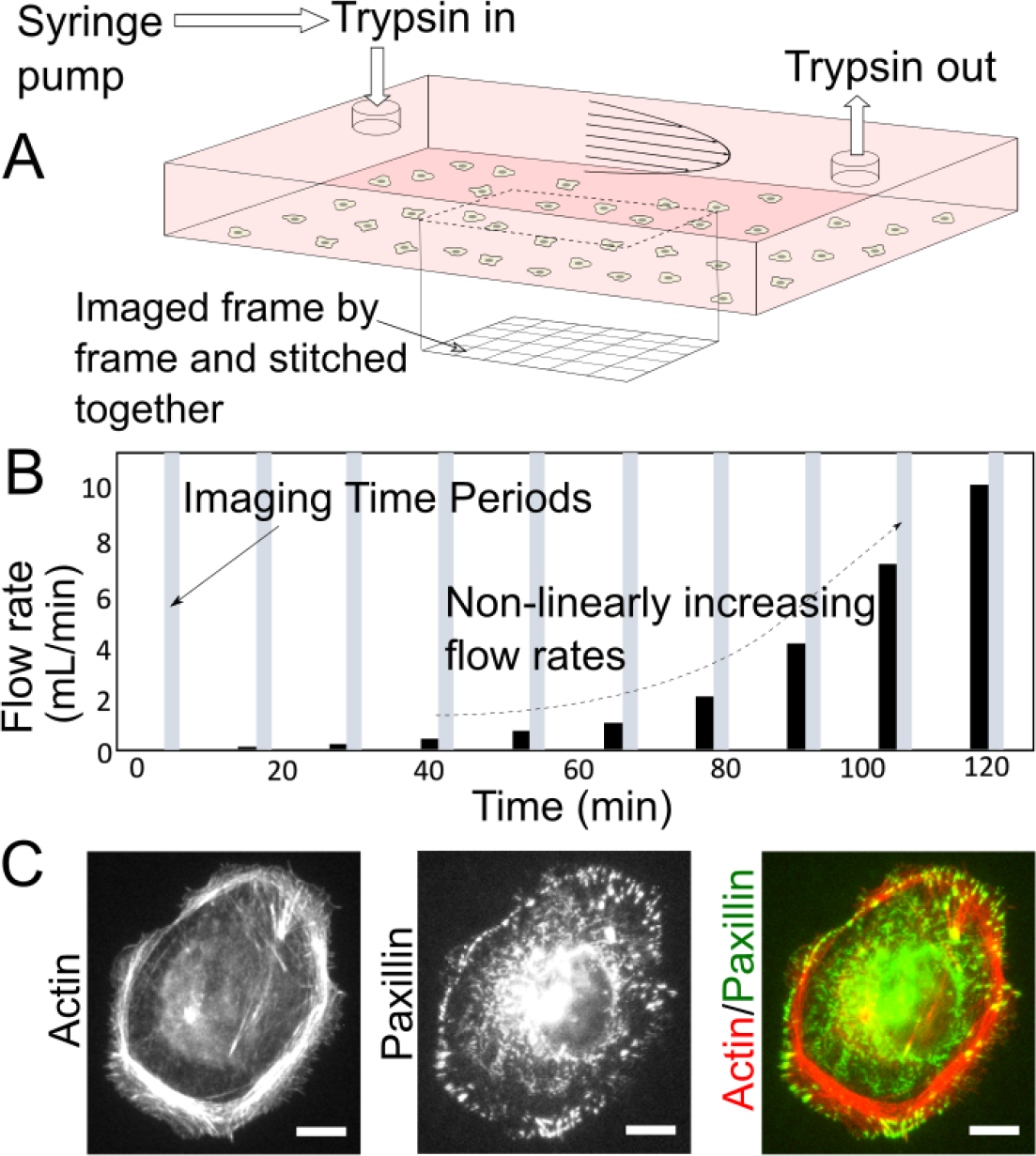
Adhesion strength assay integrating shear flow and trypsin treatment for strongly adherent cells. (A) Schematic illustration of the microfluidic chamber with the inner bottom surface coated with collagen I and plated with cells. A trypsin solution (with 0.25% trypsin and 0.1% EDTA) is flown through with a syringe pump at flowrates specified in (B). The expected parabolic velocity profile for laminar flow is shown schematically. The region over which the cells are imaged is shown with a dotted line. Imaging is carried out frame by frame over the shown region and then stitched into a composite image. (B) Temporal layout of trypsin flows (at the flowrates indicated on the y-axis) and intermittent imaging periods. (C) Immunofluorescence image of an MDCK cell adherent on collagen I coated glass. The actin cytoskeleton and paxillin-marked focal adhesions are shown. Scale bar is 10 μm.

[44] which is equivalent to a shear stress τ given by

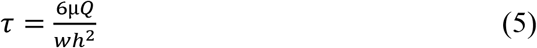

Thus, for volumetric flowrates in the range 0.1 to 10 mL/min, the corresponding shear stress at the bottom wall of our channel was in the range 0.07 to 6.7 Pa.

Cells used in our assay were adherent on the collagen I-coated bottom glass surface within the chamber and adhere via integrin receptors that form micron-scale focal adhesions that are coupled to the actin cytoskeleton within the cell (Fig. 2C shows an immunofluorescence staining of focal adhesions and actin). In order to achieve high throughput, we imaged nearly 1 cm^2^ of the surface by acquiring a grid of overlapping images (Fig. 2A). We obtained cell phase images to assess gross cell morphology and the extent of cell spreading (Fig. 3A). We also obtained nucleus fluorescence images (using a nuclear marker, Fig. 3A) to easily locate cells and ensure that single cells identified as such using phase imaging are indeed single. Finally, imaging of the plasma membrane of the cells was enabled by a plasma membrane marker (Fig 3A) and aided in cell shape segmentation and analysis (Fig. 3B). While the phase and plasma membrane images were taken just at the initial timepoint for subsequent cell morphology analysis, the nucleus images were taken at all time points and enabled the identification of the flow rate (Fig. 2B) that dislodged each identified cell. Thus, the shear stress that dislodged each cell, in combination with simultaneous trypsin treatment, was a quantitative read out of the adhesion strength of the cells.

**Figure 3.**
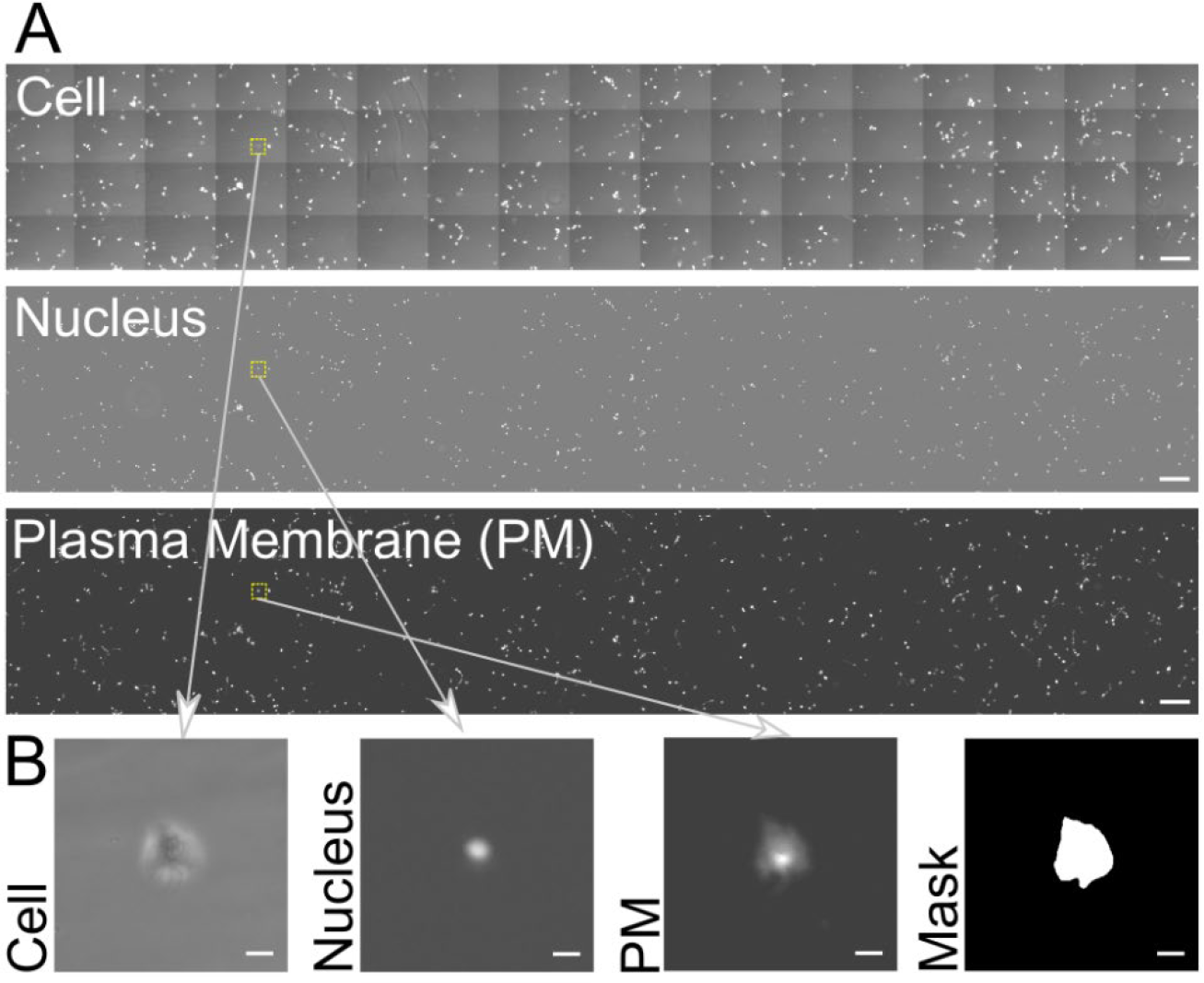
(A) Phase image of cells, fluorescence image of nuclei and fluorescence image of plasma membrane from part of the imaged region, acquired at the beginning of the assay. Scale bar is 600 μm. The small yellow dotted rectangles indicate the region shown in (B). (B) Cell, nuclear and plasma membrane images of a single cell. The corresponding cell binary mask is also shown. Scale bar is 40 μm. Note that the actual acquired image is larger, about three times as large as that shown in (A).

Morphological analysis of each cell (as adherent on collage I coated glass at the initial timepoint, before trypsin treatment and shear flow) yielded its cell area as well as circularity. The distribution of the cell area and circularity of all cells assessed in our assay are shown in Fig. 4. These distributions revealed a few aspects of the cells that were considered for adhesion strength assessment. Most notable is the lack of any overall correlation between cell area and circularity in this cell type (MDCK). However, for very small cell areas of a couple of hundred μm^2^ or less, the circularity is, on average, close to 1. This is indicative of cells that largely stayed rounded, i.e., those that adhered, but did not spread well over the surface. In fact, a circularity close to 1 is most frequent among all the cells, as evident from the histogram of circularity. The cell area varies over a wide range, from about 100 μm^2^ to under 5000 μm^2^. The majority are less than 1000 μm^2^ in cell area, but a significant fraction of cells are still above 1000 μm^2^. When cells with this distribution of shape characteristics were subject to simultaneous trypsin treatment and shear flow, the percentage of cells that survived (remained adhered) after each shear stress is plotted in Fig. 5A. Based on the cell area and circularity characteristics of the cells that survived each shear stress, the different shear stress values used in our method clustered into three categories – low: <= 0.3 Pa, medium: 0.3 to 2.7 Pa and high: >= 2.7 Pa (dashed ovals in Fig. 5A).

**Figure 4.**
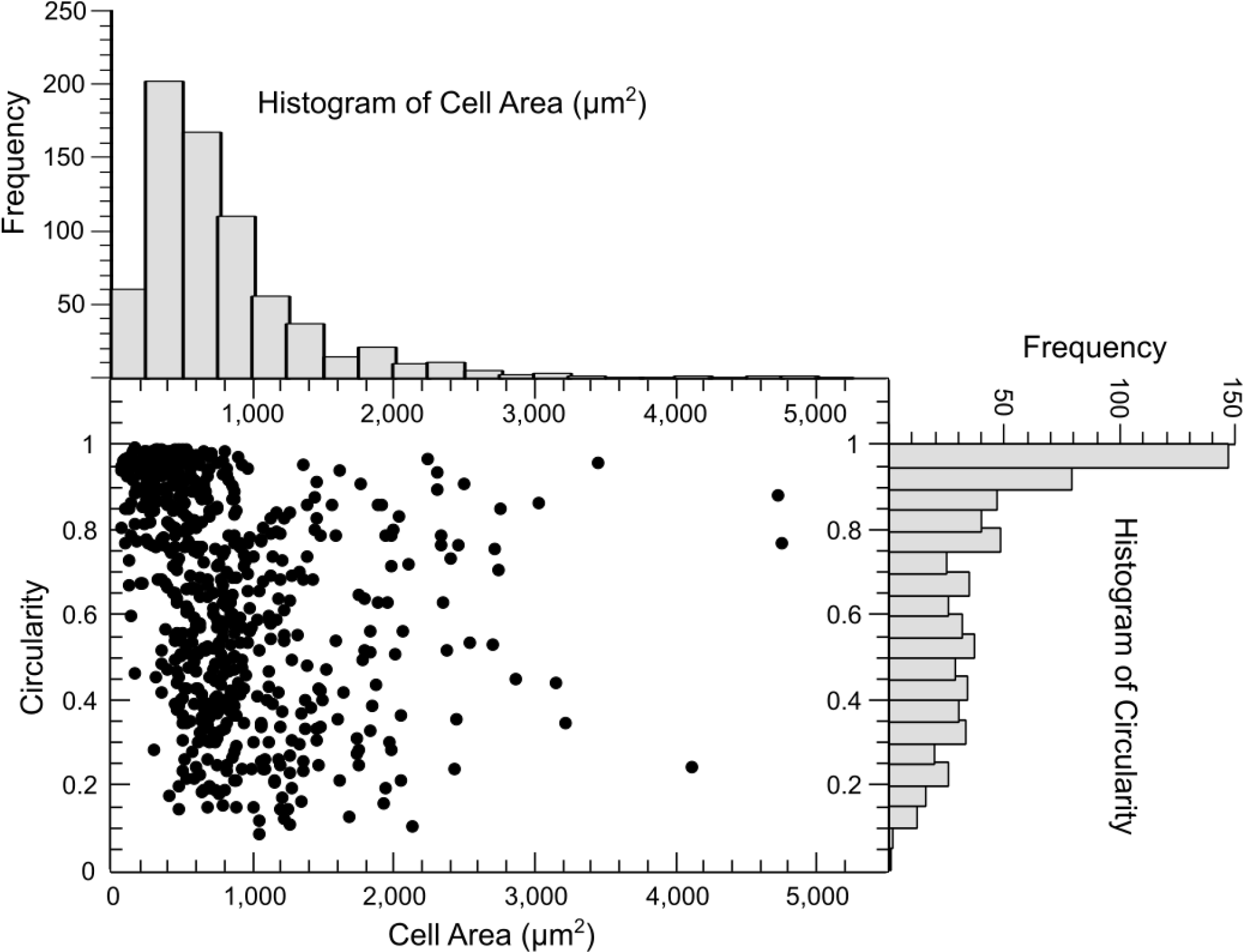
Morphological characteristics of all the cells considered in our assay. Plot of the circularity of cells versus their cell area (at the beginning of the assay) is shown. Corresponding histograms of the cell area and circularity are shown in the top and right, respectively.

**Figure 5.**
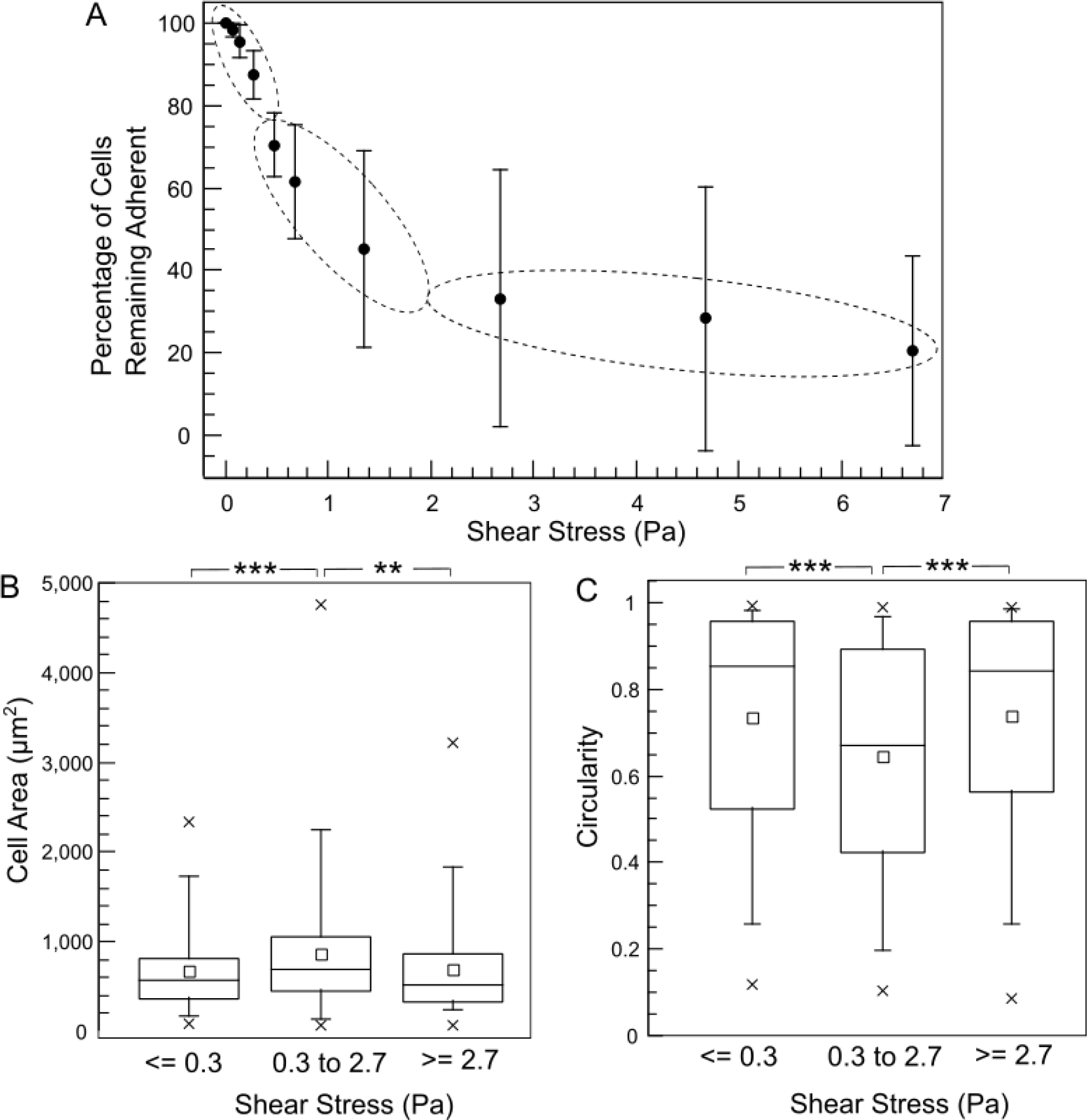
(A) Plot of the percentage of cells still adherent versus the shear stress applied by the flow of the trypsin solution. Each dot represents the mean for a given shear stress and error bars are ± standard deviation. The shear stresses are grouped into low, medium and high categories as shown by the dotted ellipses, for further analysis in (B, C). (B) Boxplot of the distribution of cell areas of cells removed by the trypsin solution with flow shear stress <=0.3 Pa, between 0.3 and 2.7 Pa, and >=2.7 Pa. (C) Boxplot of the distribution of circularity of cells removed by the trypsin solution with flow shear stress <=0.3 Pa, between 0.3 and 2.7 Pa, and >=2.7 Pa. For the boxplots in (B) and (C), the small squares represent the mean, the horizontal line represents the median, the box represents the 25^th^ to 75^th^ percentile, whiskers represent the 5^th^ and 95^th^ percentiles, and the crosses represent the minimum and maximum values. A total of 699 cells pooled from 3 experiments were considered.

Fig. 5B shows the distribution of cell areas of the cells that came off the surface at each of the shear stress categories (combined with trypsin treatment). As expected, the cell areas of the cells that come off the surface at the low shear stress category are lower, on average, than those that come off at the medium shear stress. This can be attributed to the rounded, weakly bound cells, facing relatively more hydrodynamic drag due to their shape, coming off readily at the lower flow rates. It is also likely that the larger cells, which can be expected to have greater adhesion strength, come off at medium rather than low shear stress. Interestingly, the cells that come off in the high shear stress category have a lower mean cell area than those that come off at the medium shear stress category. This seems counterintuitive, but can be understood, by taking note of the circularity of cells that came off in each shear stress category (Fig. 5C). The circularity of the cells that came off at the high shear stress is higher, on average, than those that came off at the medium shear stress category. The rate at which trypsin will diffuse to the adhesion receptors beneath the central region of the cells with a higher circularity can be expected to be lower than the rate at which it can reach the adhesion receptors underneath cells with low circularity. This factor is a likely explanation for why cells with smaller cell area but larger circularity, on average, come off in the high shear stress category compared to the medium shear stress category. Furthermore, to illustrate the insights obtained by considering morphological data, segmented cell boundaries of cells with maximum, mean and minimum cell area as well as those with maximum, mean and minimum cell circularity for each of the shear stress categories are shown in Fig. 6. Cells with the least area in each category are rounded cells (Fig. 6, as evident from the near circular shape of the minimum cell area cases for all shear stress categories). It is also evident that cells with long processes, even if the processes may be adhesive only at their ends, are those that possess least circularity when analyzed using 2D morphology. Fig. 6 also shows that the low shear stress category predominantly removes rounded cells, since the cell with mean circularity is itself rather rounded. Finally, Fig. 6 also illustrates that cell area or circularity by itself is a poor predictor of the adhesion strength – i.e., cell heterogeneity in adhesion strength is high for a given morphological profile. What role surface protein coating and cell type [34] play in this ought to be investigated further.

**Figure 6.**
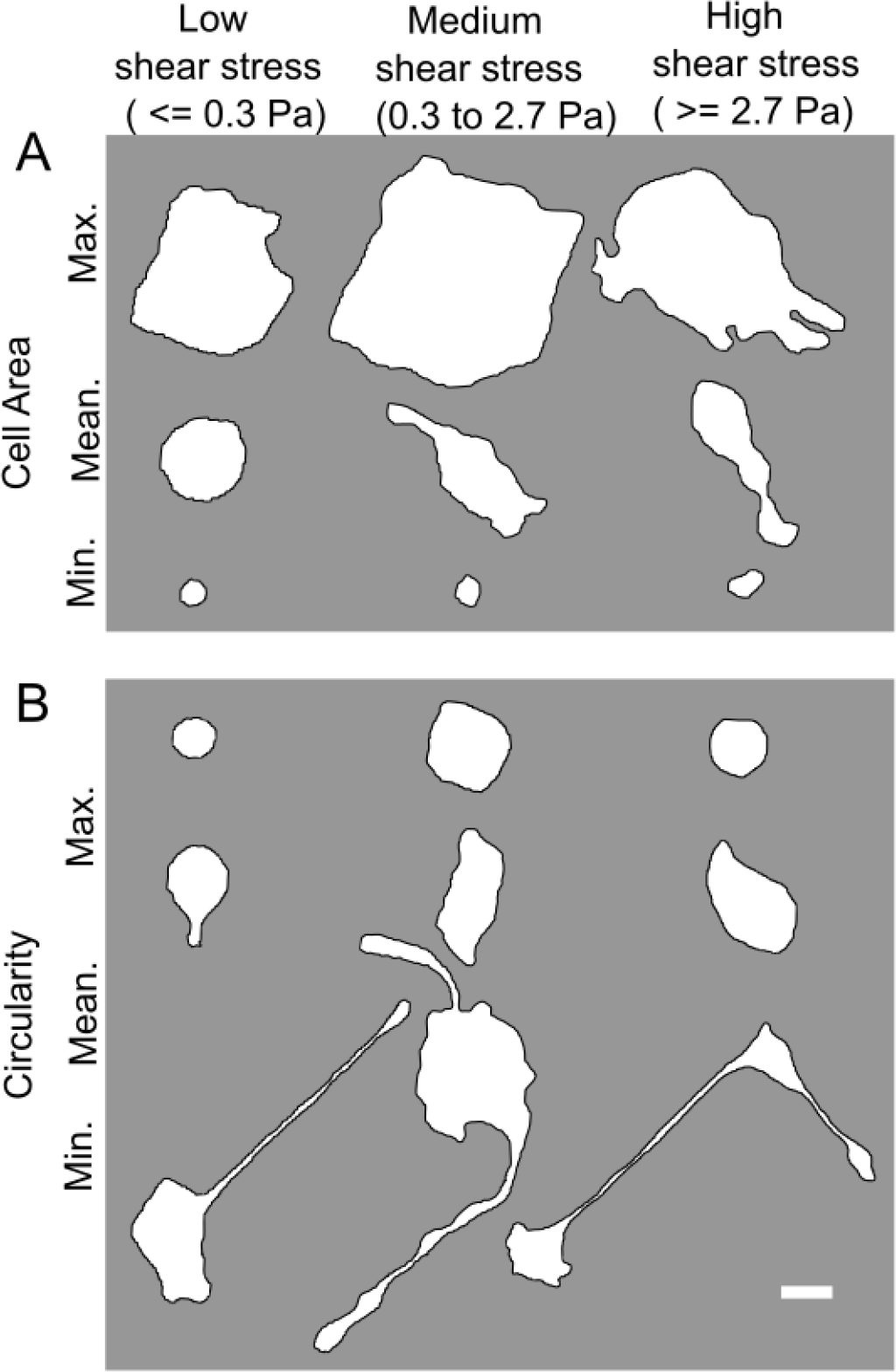
(A) Segmented cell boundaries of cells with the maximum, mean and minimum cell areas among the cells that were dislodged in the low, medium or high shear stress categories, in addition to concomitant trypsin treatment. (B) Segmented cell boundaries of cells with the maximum, mean and minimum cell circularity among the cells that were dislodged in the low, medium or high shear stress categories, in addition to concomitant trypsin treatment. Scale bar at the bottom right is 20 μm.

In combination with trypsin treatment, the range of wall shear stress employed in our method is of two orders of magnitude and is therefore applicable for cell samples that have a wide range of cell adhesion strengths. Trypsin treatment and shear flow also act synergistically to aid in the detachment of even strongly adhesive cells. That is, trypsin treatment not only leads to progressive tuning down of the adhesions, but also concomitant rounding of the cells which in turn increases the hydrodynamic shear forces experienced by the cells. Furthermore, the dynamics of deadhesion via trypsin also depends on the contractility of cells [45]. This implies that highly contractile cells will likely round up more as trypsin treatment progresses and consequently be dislodged by hydrodynamic shear forces. Thus, the relative dynamics of cell deadhesion by trypsin treatment and shear flow may vary from one cell type to another. Here, we assessed the adhesion strength of cells plated for 2 hours before the assay. For some surfaces, plating cells overnight may be more appropriate in order to allow cells to develop mature focal adhesions beneath the cell surface [39]. This can be addressed in our assay by plating cells overnight, but maintaining a flow of cell culture media of very low magnitude to keep the cells viable during this longer plating period before the assay begins [36]. It is worth noting that trypsin sensitivity may depend on the specific cell adhesion proteins involved [46], therefore different trypsin concentrations can also be used in the assay to match it to the adhesion strength of a particular cell type. In fact, we found that MDCK cells on collagen I coated glass were not dislodged by the shear stresses we employed when the buffer used contained 0.5 mM EDTA but not trypsin and only a minority of the cells were dislodged by 0.05% trypsin (data not shown).

## Conclusion

We have proposed and demonstrated an assay for adhesion strength assessment that is high throughput, widely accessible and allows cell morphological profiling. The low shear stress requirement is due to the simultaneous use of trypsin treatment and allows the use of off-the-shelf chambers to implement the assay. This also makes the assay amenable to be widely adopted. Cell morphological profiling showed that cell adhesion strength heterogeneity for a given size and shape is high. While we demonstrate its use for a strongly adherent epithelial cell line (MDCK), it can similarly be employed with other epithelial cells, endothelial cells and fibroblasts that form many large focal adhesions and adhere to the substrate strongly as well as more weakly adherent cell lines by tuning the trypsin concentration used. Promisingly, the absence of extensive surface treatment or low height microfluidic channels in our assay also means that it can be extended to use soft substrates [47, 48] atop glass to study how substrate stiffness influences the cell adhesion strength.

## Acknowledgments

V.M. acknowledges support from the National Institute of General Medical Sciences of the National Institutes of Health under award number 2R15GM116082.

## Author Declarations

### Ethics Approval

This investigation was conducted using an immortalized cell line. Therefore, no ethics approval was required.

### Author Contributions

**Antra Patel**: Methodology; Investigation; Writing/Original Draft Preparation. **Bhavana Bhavanam**: Investigation. **Trevor Keenan**: Methodology. **Venkat Maruthamuthu**: Conceptualization; Visualization; Project Administration; Writing/Review & Editing; Funding Acquisition

### Data Availability Statement

The data that support the findings of this study are available from the corresponding author upon reasonable request.

